# Neurotransmitter imbalance in the brain and Alzheimer’s pathology

**DOI:** 10.1101/220699

**Authors:** Stuart G. Snowden, Amera A. Ebshiana, Abdul Hye, Olga Pletnikova, Richard O’Brien, An Yang, John Troncoso, Cristina Legido-Quigley, Madhav Thambisetty

## Abstract

**INTRODUCTION:** Three of the four treatments for Alzheimer’s disease are cholinesterase inhibitors targeting the pathological reduction of acetylcholine levels. Here we aimed to determine the role of other neurotransmitter pathways in AD pathology.

**METHODS:** Tissue samples were obtained from three groups, controls, AD and ‘asymptomatic AD’ i.e. cognitively normal individuals that had significant AD neuropathology. Three brain areas were studied, the middle frontal gyrus (MFG) the inferior temporal gyrus (ITG) and the cerebellum.

**RESULTS:** 11 of 15 measured metabolites were shown to be associated with disease. Decreases in dopamine were seen in the ASYMAD group in the MFG when compared to control and AD patients (FC=0.78, p=4.1×10^-3^). In AD patients changes were mainly seen in the ITG’s inhibitory GABAergic system.

**DISCUSSION:** These results indicate that dopamine could be depleted in brains with Alzheimer’s pathology but intact cognition, while and imbalance of several neurotransmitters is evident in the brain of AD patients.

## 1. Introduction

Dementia is a devastating illness for both patients and their families, with Alzheimer’s disease (AD) estimated to account for up to 80% of total dementia cases. The ‘World Alzheimer’s report 2015’ estimates that there are approximately 46 million AD patients worldwide, with this number expected to rise to over 130 million by the middle of the century (1). As well as a significant human cost, AD also represents a major financial burden with worldwide costs related to AD expected to reach $1 trillion dollars in 2018 (1).

Cholinesterase inhibitors make up three of the four approved AD treatments (Donepezil, Rivastigmine and Galantamine) making inhibition of acetylcholinesterase the leading therapeutic strategy for the treatment of AD symptoms (2, 3). There is a significant body of literature that has suggested that the cognitive deficits associated with Alzheimer’s disease are the result of lower levels of acetylcholine in the brain resulting from dysfunction of cholinergic neurons (4-6). The role of non-cholinergic neurotransmitter systems in AD pathogenesis has received less attention. While levels of non-cholinergic neurotransmitters in the brain have been associated with Alzheimer’s pathology (7-11), their role in mediating the onset of symptoms is less well understood. In this study, we analysed data from non-targeted metabolomics to compare differences in neurotransmitters and neurotransmitter-associated metabolite levels in brain tissue samples from the autopsy cohort of the Baltimore Longitudinal Study of Aging (BLSA). We studied three groups of BLSA participants, AD patients, cognitively normal controls and ‘asymptomatic AD’ (ASYMAD; i.e. individuals with significant AD neuropathology at death but with no evidence of cognitive impairment during life). We studied three distinct brain regions in these individuals that are differentially effected by core pathological features of AD, the inferior temporal gyrus that is especially vulnerable to neurofibrillary tau tangles, the middle frontal gyrus which is susceptible to the accumulation amyloid plaques and the cerebellum which is resistant to classical AD pathology (12). Our aim was to test associations between AD neuropathology and metabolism in a variety of neurotransmitter systems.

## 2. Methods

### 2.1 Sample Information

The BLSA is a prospective, ongoing cohort study of community-dwelling volunteer participants in Baltimore begun in 1958. As such, it is among the largest and longest-running longitudinal studies of aging in the United States (13, 14). In general, at the time of entry into the study, participants had no physical or cognitive impairment. Detailed examinations, including neuropsychological assessments and neurological, laboratory, and radiological evaluations, were conducted every 2 years. Since 2003, participants older than 80 years have received yearly assessments. Written informed consent was obtained at each visit, and the study was approved by the local Institutional Review Board and the National Institute on Aging. After each visit, cognitive status was considered at consensus diagnosis conferences relying on information from neuropsychological tests as well as clinical data as described previously (15). Diagnoses of dementia and Alzheimer’s disease (AD) were based on DSM-III-R (16) and the NINCDS-ADRDA criteria (17) respectively.

Brain tissue samples were collected through the autopsy sample of the BLSA. The autopsy program of the BLSA was initiated in 1986. We have previously described the study protocol in detail. Briefly, the mean age at death in the autopsy sample is 88.3 ± 7.3 years (range 69.3-103.2), and the mean interval between last evaluation and death is 8.7± 6.7 months (18). As reported previously, the autopsy subsample is not significantly different from the BLSA cohort as a whole in terms of the rates of dementia and clinical stroke (19). Table 1 describes the demographic characteristics of the participants whose brain tissue samples were used in this study.

**Table 1.** Clinical Characteristics of study participants.

|  | **Control** | **Asymptomatic** | **Demented** |
| --- | --- | --- | --- |
| Participants (f/m) | 14 (4/10) | 15 (5/10) | 14 (7/7) |
| Age at death (Years)* | 82.6 ± 11.0 (64.2-99.2) | 89.2 ± 7.9 (71.9-96.4) | 87.9 ± 8.9 (62.9-98.7) |
| MMSE<sup>†</sup> | 27.8 ± 2.4 | 29.0 ± 0.9* | 23.0 ± 6.9** |
| PMI (Hours)<sup>‡</sup> | 16.9 ± 6.4 (7.0-28.0) | 14.8 ± 8.1 (2.0-33.0) | 14.7 ± 6.0 (3.0-23.0) |
| Cholinesterase/NMDA agonist usage | 0/0 | 0/0 | 2/0 |

\* values are reported as the mean ± standard deviation, and range, <sup>†</sup> values are reported as the mean ± standard deviation <sup>‡</sup> values are reported as the mean ± standard deviation, and range. MMSE; Mini-mental state examination, NMDA; N-methyl-D-aspartate, PMI; post mortem interval.

### 2.2 Data acquisition

The majority of the data described in this paper were generated in a previously published study with a detailed description of the acquisition, analysis and annotation of thirteen metabolites: tyrosine, L-DOPA, dopamine, aminobutanal, arginine, aspartate, GABA, glutamate, glutamine, guanidinobutanoate, glycine, guanosine and ornithine (20). Additionally new data concerning measures of serotonin and tryptophan were acquired using the Biocrates platform. To extract metabolites from brain tissue, samples were homogenized using Precellys® with ethanol phosphate buffer. Samples were then centrifuged and the supernatant was used for analysis. The fully automated assay was based on liquid chromatography-tandem mass spectrometry (LC-MS/MS; amino acids) using a SCIEX 4000 QTrap® mass spectrometer (SCIEX, Darmstadt, Germany) with electrospray ionization. Brain tissue concentration was absolute concentration expressed as nmol/mg tissue

### 2.3 Statistical methods and pathway mapping

To compare the abundance of neurotransmitters and their associated metabolites among 3 groups (CN, ASYMAD and AD), we used Mann–Whitney U test for pairwise comparisons. To control for type 1 errors in the p-values calculated using the Mann-Whitney U test a Benjamini-Hochberg procedure performed in ‘R’ with the results reported in Table 2. The relationship of metabolite abundance to measures of neuritic plaque and neurofibrillary tangle burdens in the brain as described by CERAD and Braak scores respectively were determined by calculating the Pearson’s product-moment correlation coefficient.

**Table 2.**
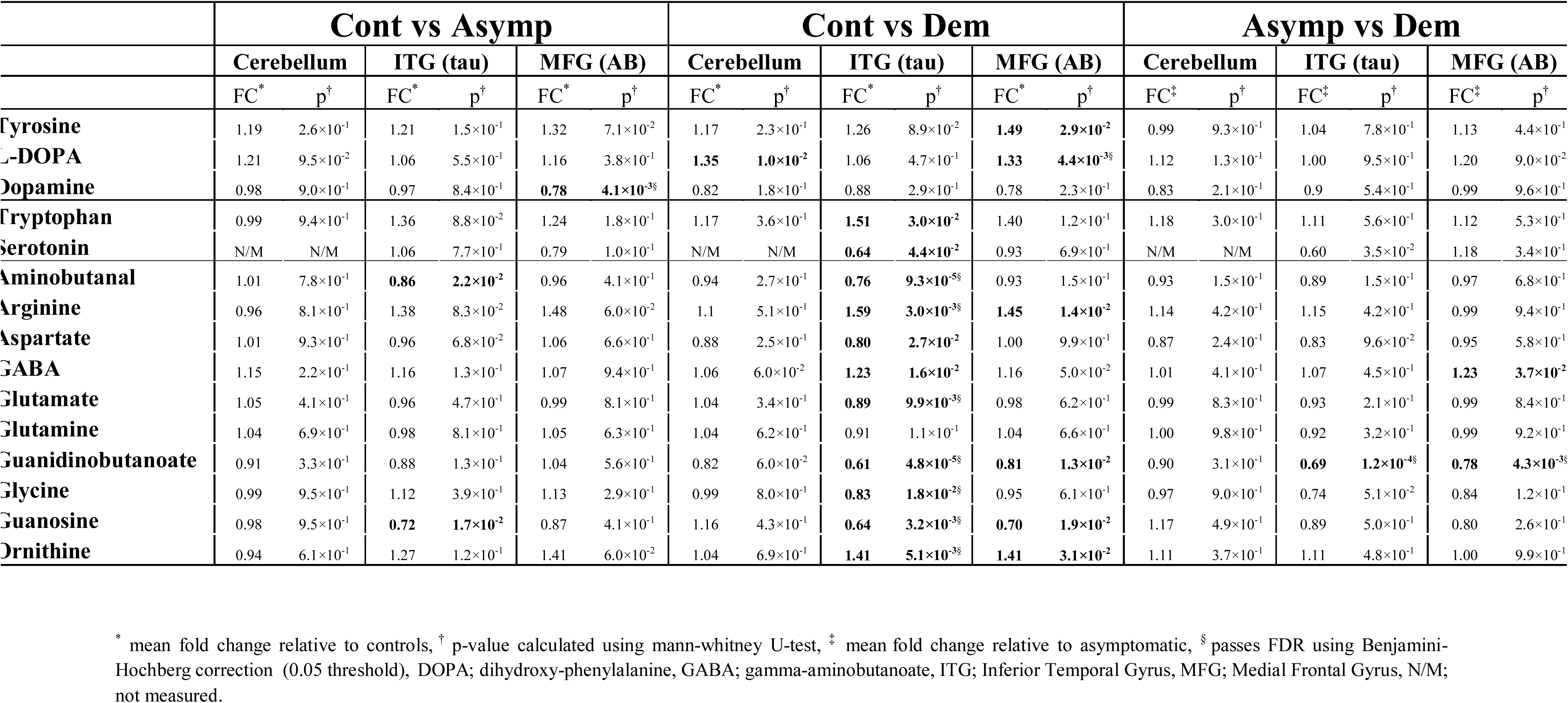
Relative changes in abundance of 15 metabolites associated with neurotransmitter metabolism between all three diagnostic groups in individual brain regions.

Pathway mapping was performed in Cytoscape v3.4.0 the architecture was determined by metabolic interactions defined in the Kyoto Encyclopaedia of Genes and Genomes (KEGG). Within the network node size is directly proportional to the fold change in metabolite abundance, with edge thickness directly proportional to the partial correlation of the two nodes it is connecting.

## 3. Results

### 3.1 Region specific analysis in the ASYMAD versus control group

The abundance of all fifteen metabolites were compared between control and ASYMAD groups (Figure 1, Table 2, Supp Figure 1). The only neurotransmitter that had an altered abundance was dopamine which was decreased in the MFG (FC=0.78, p=4.1×10^−3^). In the ITG aminobutanal and guanosine which are both involved in the metabolism of neurotransmitters were also decreased (FC=0.86, p=2.2×10^−2^ and FC=0.72, p=1.7×10^−2^ respectively) (Figure 1, Table 2, Supp Figure 1).

**Figure 1.**
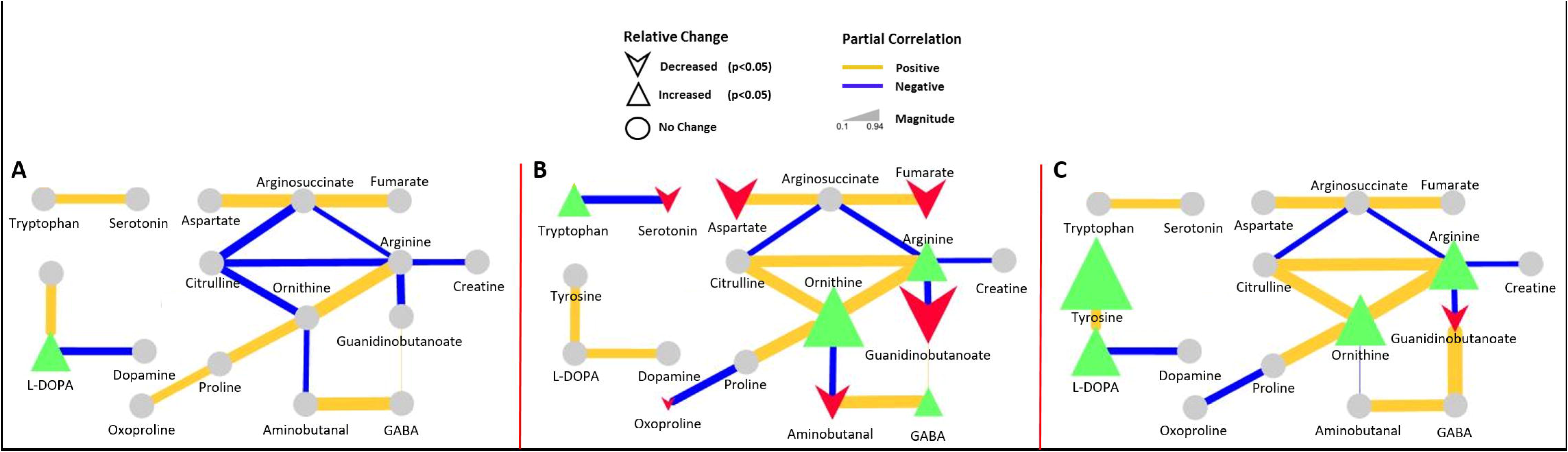
Showing pathway analysis of the association of neurotransmitter metabolism to Alzheimer’s disease in human brain. Metabolites significantly increased in abundance (p<0.05) and shown as green triangles and metabolites significantly decreased in abundance (p<0.05) and shown as red chevrons with the size representing the magnitude of the change. Grey circles represent metabolites that were not significantly associated with disease. A) shifts observed in the cerebellum, B) shifts observed in the inferior temporal gyrus, C) shifts observed in the middle frontal gyrus.

### 3.2 Region specific analysis in the AD versus control group

In the comparison of control versus AD groups, changes were observed mainly in the ITG. In the ITG excitatory neurotransmitters glutamate and aspartate exhibited a lower abundance (p<0.05 & FC=0.8) in AD patients. Also in the ITG, inhibitory neurotransmitters glycine, and serotonin were decreased whilst GABA was increased (p<0.05, FC=0.8 and 1.2). A number of neurotransmitter precursors were also increased: ornithine, arginine and tryptophan (all p<0.05) whilst guanidobutanoate, guanosine, aminobutanal were all significantly decreased (all p<0.05) in the ITG of AD patients (Figure 1, Table 2, Supp Figure 1). In the MFG dopamine precursors L-DOPA and tyrosine were the only metabolites to be increased with disease. A significant increase in L-DOPA was also the only significant difference observed in the cerebellum. (Figure 1, Table 2, Supp Figure 1).

### 3.3 Region specific analysis in the AD versus ASYMAD groups

In the comparison of AD versus ASYMAD, two changes were observed in the MFG: GABA was increased (FC=1.23, p=3.7×10^-2^) and guanidobutanoate was decreased (FC=0.78, p=4.3×10^-3^). In the ITG, guanidobutanoate was also decreased (FC=0.69, p=1.2×10^-4^) (Figure 1, Table 2, Supp Figure 1).

### 3.4 Correlation of metabolite abundance and measures of Alzheimer’s pathology

When examining the relationship between metabolite abundance and measures of pathology and cognitive performance several weak but significant correlations were observed. Of the 15 measured metabolites, all (with the exception of dopamine, glutamate and glutamine) correlated with Braak and CERAD scores in all regions (r^2^> 0.2, p<0.05) (Supplemental Table 1). Correlation analysis to investigate the relationship between metabolite abundance and cognitive performance showed that arginine, aspartate, aminobutanal and guanidobutanoate correlated with MMSE (r^2^> 0.2, p<0.05) in all regions (Supplemental Table 2), GABA, aspartate, tyrosine, DOPA, ornithine, arginine, guanidobutanoate and aminobutanal correlated with Benton’s visual retention index (r^2^> 0.2, p< 0.05) (Supplemental Table 2) and guanidobutanoate correlated with the Boston naming score. Running Title: Neurotransmitter metabolism in Alzheimer’s brain

## 4. Discussion

The metabolism of neurotransmitters is an important consideration in the pathology of all neurological diseases. In this study we measured the metabolism of three key excitatory neurotransmitters dopamine, glutamate, and aspartate, as well as three inhibitory neurotransmitters, serotonin, glycine and GABA. We tested to see if any observed modifications in neurotransmitter pathways were associated with the asymptomatic AD group. After this we wanted to determine if specific brain regions exhibited unique differences in neurotransmitter metabolism. Considering the pathways studied here, the dopaminergic pathway, which was depleted in the asymptomatic patients, was most strongly associated with amyloid and tau burden in the MFG.

### 4.1 Dopaminergic depletion in brains with neuropathology and normal cognition

Dopamine is a catecholamine neurotransmitter (21, 22) which plays several important roles in the brain acting via 4 distinct pathways, the mesolimbic, mesocortical, nigrostatial and tuberoinfundibular pathways. These pathways are responsible for regulating mood, and aiding cognitive and motor function. Impairment of this system potentially causes depression (23), memory loss (24) and impaired motor control observed in patients with Alzheimer’s disease.

Dopamine does not cross the blood brain barrier and is synthesised in two steps from the essential amino acid tyrosine, with the initial conversion of tyrosine to L-DOPA catalysed by tyrosine hydroxylase (TH) with the subsequent conversion of L-DOPA to dopamine catalysed by aromatic amino acid decarboxylase (AAAD). The greatest reduction in dopamine in the MFG was observed in the asymptomatic patients (Figure 2, Supp Figure 1). In the MFG this pathway shows an increase in the abundance of tyrosine and L-DOPA, the precursors of dopamine, followed by a reduction in the abundance of dopamine, suggesting a decrease in the abundance or activity of both TH and AAAD.

**Figure 2.**
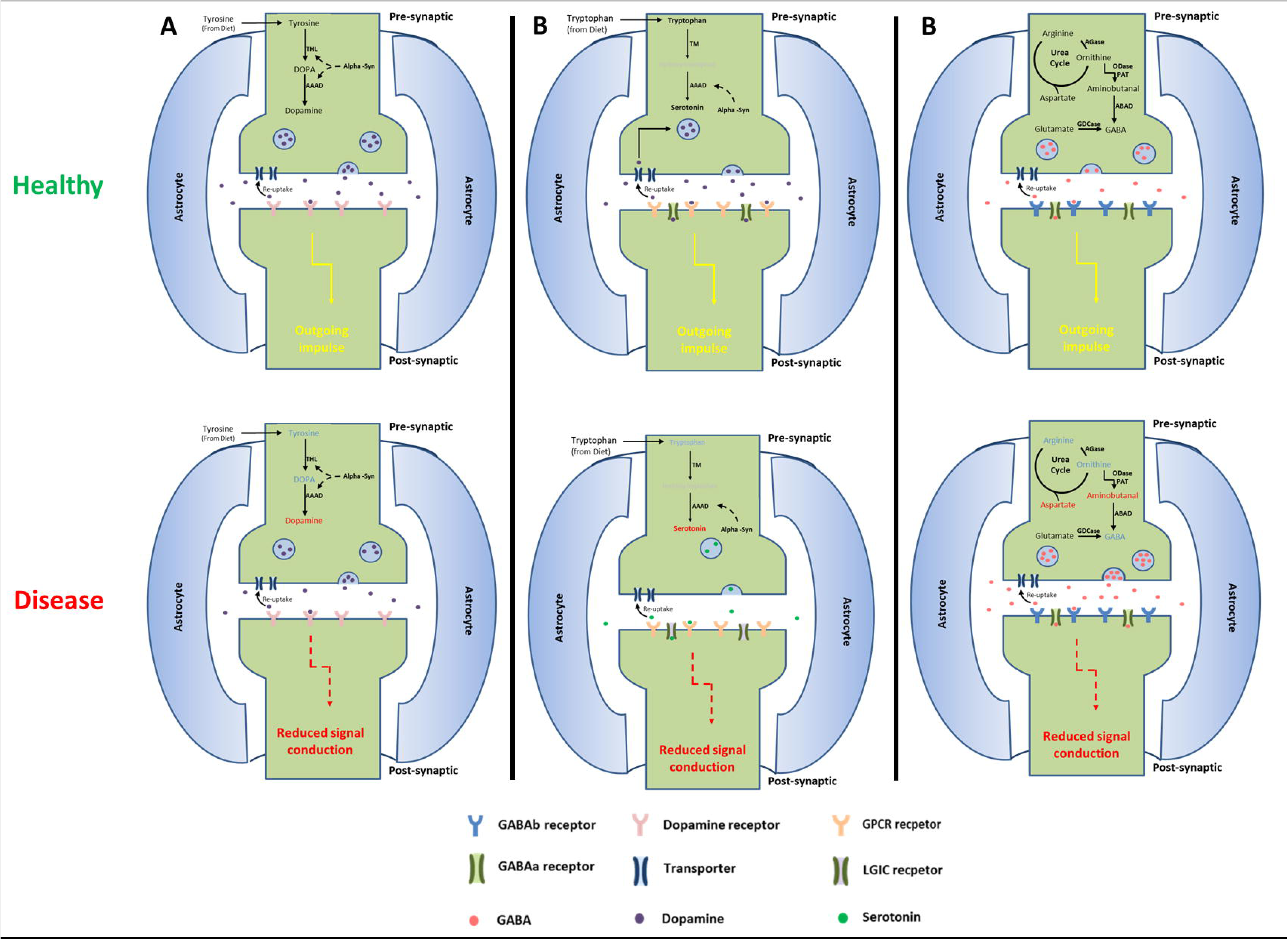
Potential modifications to synaptic transmission based on alterations in metabolism associated with Alzheimer’s pathogenesis. (A) Effects of decreased dopamine synthesis on synaptic transmission. (B) Effects of decreased serotonin synthesis on synaptic transmission. (C) Effects of disrupted GABA metabolism on synaptic transmission. Metabolites or processes shown in **red** are decreased in abundance, **blue** are increased, **black** were measured but unchanged, and grey were non measured intermediates, solid lines represent metabolic reactions and dashed lines represent protein protein interactions. AAAD: aromatic amino acid decarboxylase, ABAD: aminobuteraldehyde dehydrogenase, AGase: arginase, alpha-syn: alpha-synuclein, GDCase: glutamate decarboxylase, ODase: ornithine decarboxylase, PAT: putrescine aminotransferase, THL: tyrosine hydroxylase, TM: tryptophan monooxygenase.

The activity of both TH and AAAD are inhibited by alpha-synuclein (25), a protein that has been shown to possess important pathological roles in a range of neurodegenerative diseases including Parkinson’s (26), Alzheimer’s (27) and Lewy bodies dementia (26-28). Studies have also shown that soluble intra-neuronal alpha-synuclein, in the absence of Lewy body pathology is increased in abundance by up to two-fold in the brain of AD patients (29). This suggests a plausible molecular mechanism by which alpha-synuclein may modulate brain dopamine concentrations in AD. Reduced dopamine could reduce the amount of the neurotransmitter released in to the synaptic cleft during synaptic transmission leading to impaired signal transduction (Figure 2). The data (Table 2) shows that shifts in dopamine metabolism are greater in the middle frontal gyrus in the ASYMAD group, suggesting that the changes in dopamine metabolism occur before memory loss occurs.

### 4.2 Neurotransmission inhibition in the inferior temporal gyrus in AD

The inhibitory neurotransmitter serotonin and its precursor tryptophan were measured in this study and decreased in the ITG of AD patients (p<0.05); but were not significant at p<0.005 (Table 2, Supp Figure 1N and 1O). Serotonin is synthesised from the essential amino acid tryptophan by tryptophan monooxygenase (TM) and aromatic amino acid decarboxylase (AAAD). As stated above, alpha-synuclein which is increased in the brains of AD patients (29) has been shown to inhibit the action of AAAD (25) suggesting a potential mechanism for the decreasing trend in serotonin synthesis that was observed in this study. The association of serotonin to the AD brain is interesting as recent reports have shown that antidepressants such as Trazodone, a serotonin antagonist, could maintain neural integrity (31, 32).

Glutamate was also reduced in the ITG of AD patients, this was surprising as glutamate activation of N-methyl-D-aspartate (NMDA) on the post synaptic neuron and its excite-toxicity have long been implicated in the pathology of AD (33-35). The impairment of the glutaminergic system in the brain leads to impairment of a range of neurological functions, including fast excitatory neurotransmission (36), memory and learning (37) and long term potentiation (38-40). The role of glutamate in AD is well known, Memantine, an NMDA antagonist is used to treat moderate to severe AD. Memantine has been shown to have affinity for dopamine receptors as well (41).

In this study GABA is increased in abundance but no change is seen in glutamate levels apart from a modest shift observed in the ITG. Whilst increased GABA production could still be coming at least in part from glutamate, the changes observed in other metabolites associated with GABA metabolism mean that the alterations may arise from multiple sources (Figure 2). GABA is the chief inhibitory neurotransmitter in the mammalian nervous system (42, 43). It does not cross the blood brain barrier (44) and in the brain is predominantly synthesised from the non-essential amino acid glutamate by the action of glutamate decarboxylase under standard physiological conditions (45). However, GABA can be synthesised via several pathways from a selection precursors, including aminobutanal by aminobutyraldehyde dehydrogenase (46), succinate semialdehyde by aminobutyrate aminotransferase (47) and guanidinobutanoate by guanidinobutyrase (Supp Figure 2). Two alternative GABA synthetic pathways, both of which start from the urea cycle, have intermediates that are significantly reduced in abundance (Supp Figure 2) suggesting that they may play a role in the dysregulation of GABA metabolism.

Regardless of the synthetic source of the increased abundance of GABA, this combined with the reduction in glutamate in the ITG (Table 2) can produce a reduction in the glutamate/GABA ratio, leading to an inhibitory environment and a reduction in the transmission of action potentials. When GABA is released into the synaptic cleft it binds to a range of transmembrane receptors on both the pre and post-synaptic neurons leading to the opening of ion channels allowing the negatively charged chloride ions to enter and positively charged potassium ions to escape the neuron (Figure 2) (48). This shift leads to loss of the transmembrane potential and hyperpolarisation of the cell membrane, inhibiting action potentials produced by excitatory neurotransmitters like glutamate.

In conclusion, our results suggest that abnormalities in dopamine neurotransmission is observed in brains with pathology but no memory problems. Combined therapeutic approaches, especially those affecting the GABAergic and serotonergic system might be useful as adjunctive treatments in AD.

## 5. Acknowledgements

This work has been supported by grants from the Libyan Cultural attaché of Libyan embassy and the European Medical Information Framework - Alzheimer’s disease (EMIF-AD). We would also like to thank Dr Mathew Arno manager of the genomics centre for permitting us to use his Tissuelyzer. We are grateful to participants in the Baltimore Longitudinal Study of Aging for their invaluable contribution. This research was supported in part by the Intramural Research Program of the NIH, National Institute on Aging.

## SUPPLEMENTAL FIGURE LEGENDS

**Supplemental Figure 1.**
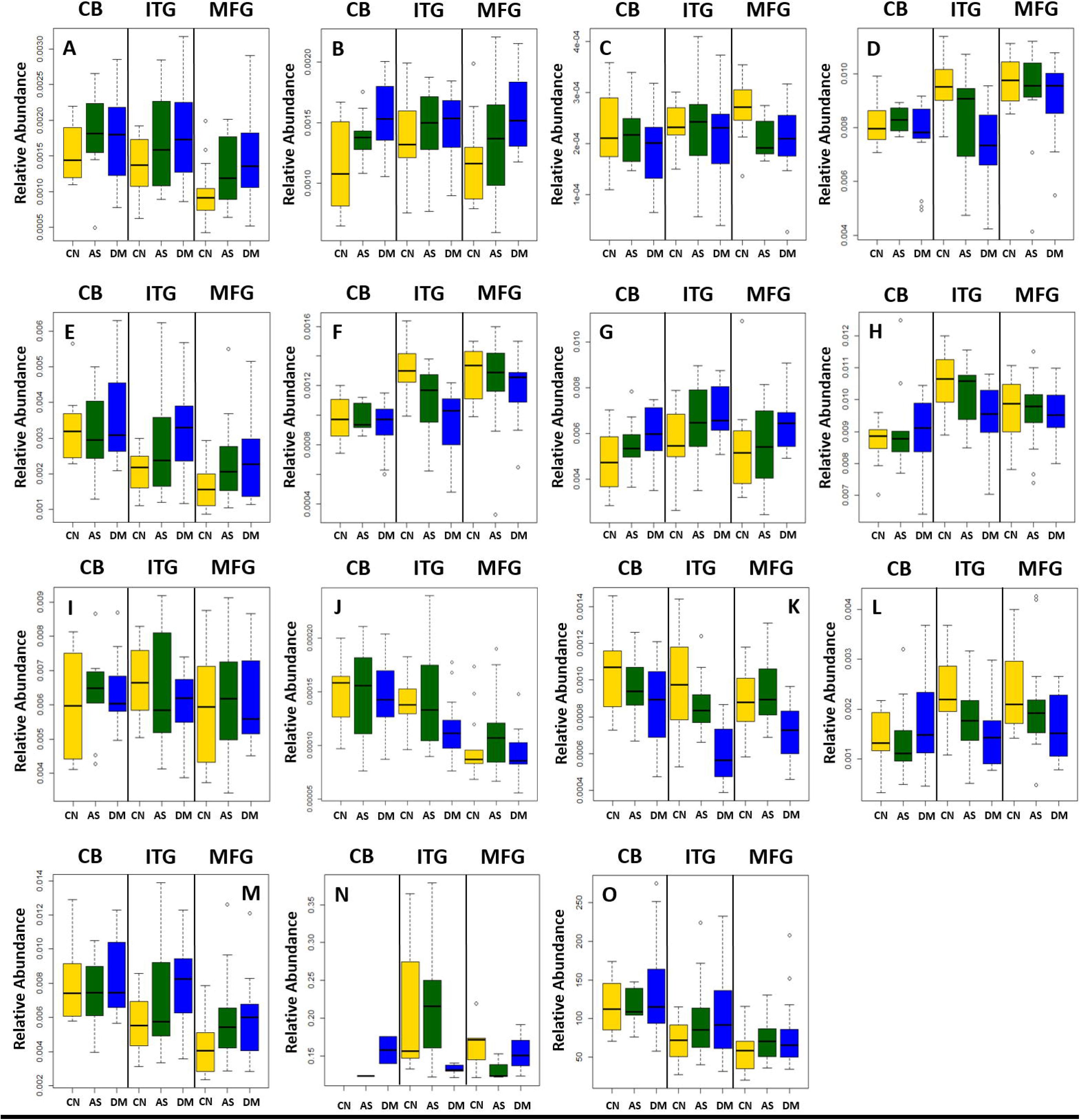
Boxplots showing the effect of disease status on the abundance of 6 neurotransmitters and 9 related metabolites in the Cerebellum, Inferior Temporal Gyrus and Middle Frontal Gyrus. Dysregulation of 5 neurotransmitters and 8 related metabolites shown by boxplots of three disease statuses separated by brain region A) tyrosine, B) DOPA C) dopamine D) aminobutanal E) arginine F) aspartate G) GABA H) glutamate I) glutamine J) glycine K) guanidinobutanoate L) guanosine M) ornithine N) serotonin O) tryptophan. DOPA; dihydroxyphenylalanine, GABA; gamma-aminobutyrate.

**Supplemental Figure 2.**
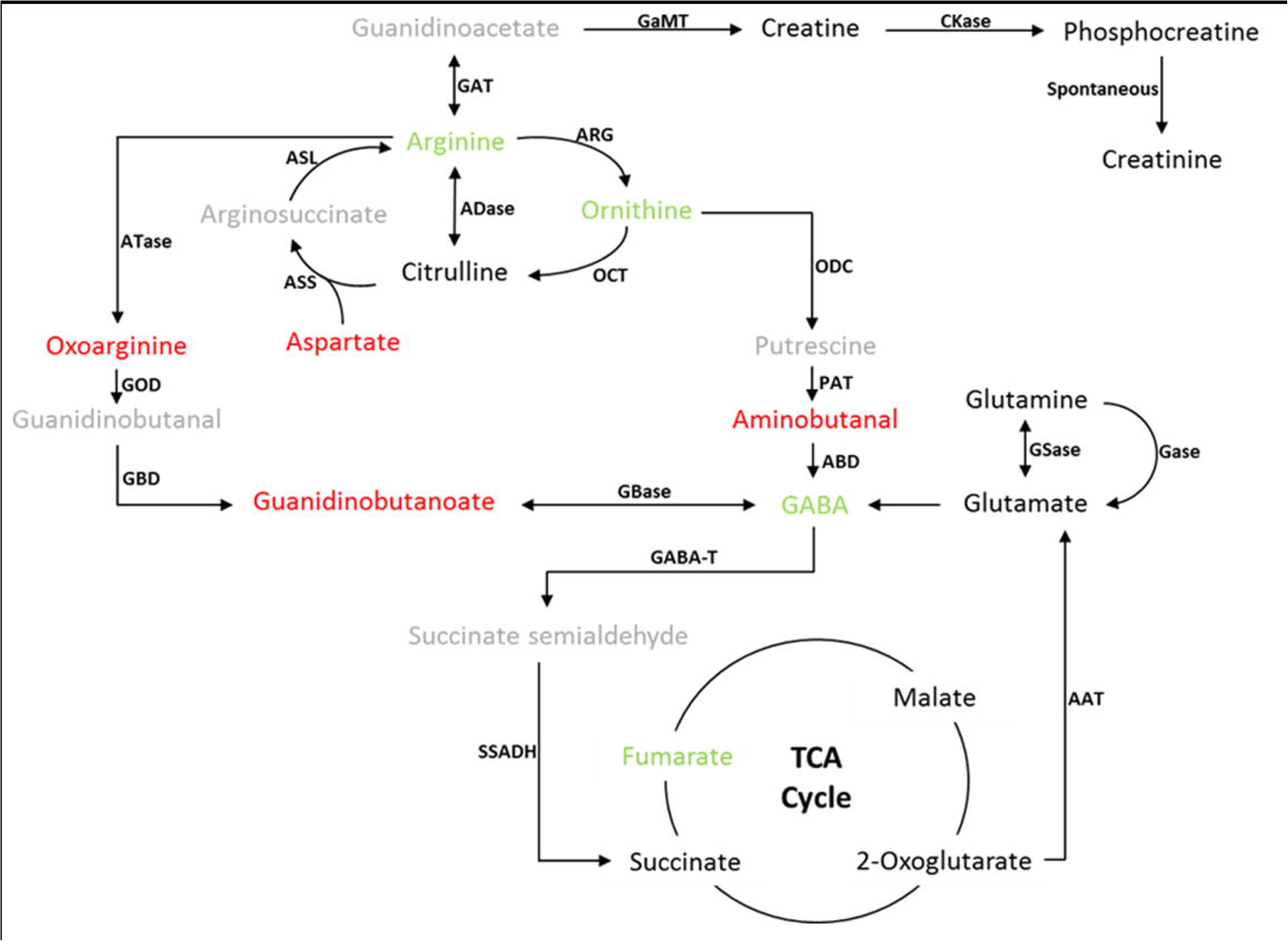
Potential metabolic modifications of arginine metabolism and GABA production related to Alzheimer’s pathogenesis. Metabolites shown in Red are decreased in abundance, blue are increased, black were measured but unchanged, and grey were non measured intermediates. ADase; Arginine deaminease, ARG; Arginase, ASL; Arginosuccinate lyase, ATase; Arginine transaminase, CKase; Creatine Kinase, GABA-T; GABA transaminase, GaMT – Guanidoacetate Methyltransferase Gase –Glutaminase GAT – Glycine amidinotransferase Gbase – Guanidobutyrase GBD – Guanidobutyraldehyde dehydrogenase GOD – Guanidino-oxopentanoate-decaroxylase GSase – Glutamine synthase OCT – Ornithine Carbomyltransferase ODC – Ornithine decarboxylase PAT – Putrescine Aminotransferase SSADH – Succinate semialdehyde dehydrogenase

**Abbreviations**
AAAD: aromatic amino acid decarboxylase
AD: Alzheimer’s disease
ASYMAD: asymptomatic AD
BLSA: Baltimore longitudinal studying of aging
CERAD: consortium to establish a registry for Alzheimer’s disease
DSM-III-R: diagnostic and statistical manual of mental disorders
GABA: gamma-aminobutyrate
ITG: inferior temporal gyrus
L-DOPA’: L-dihydroxyphenylalanine
MFG: middle frontal gyrus
NINCDS-ADRDA: national institute of neurological and communicative disorders and stroke – Alzheimer’s disease and related disorders association
NMDA: N-methyl-D-aspartate
TH: tyrosine hydrolase
TM: tryptophan monooxygenase

